# Various localized epigenetic marks predict expression across 54 samples and reveal underlying chromatin state enrichments

**DOI:** 10.1101/030478

**Authors:** Lalita Devadas, Angela Yen, Manolis Kellis

**Affiliations:** Lexington High School, Lexington, Massachusetts 02421, USA; Broad Institute of MIT and Harvard, Cambridge, Massachusetts 02142, USA; MIT Computer Science and Artificial Intelligence Laboratory, Cambridge, Massachusetts 02139, USA

## Abstract

Here, we predict gene expression from epigenetic features based on public data available through the Epigenome Roadmap Project [1]. This rich new dataset includes samples from primary tissues, which to our knowledge have not previously been studied in this context. Specifically, we used computational machine learning algorithms on five histone modifications to predict gene expression in a variety of samples. Our models reveal a high predictive accuracy, especially in cell cultures, with predictive ability dependent on sample type and anatomy. The relative importance of each histone mark feature varied across samples. We localized each histone mark signal to its relevant region, revealing that chromatin state enrichment varies greatly between histone marks. Our results provide several novel insights into epigenetic regulation of transcription in new contexts.

## Introduction

Epigenetic regulators of gene expression (the process that converts genetic information into gene product like mRNA and eventually proteins) include chromatin features, which alter how accessible the area surrounding DNA is to transcriptory enzymes [2]. Histones (proteins which allow for compaction of DNA) can be chemically modified to change the structure of chromatin [3]. Histone modifications like methylation, acetylation, and phosphorylation have been shown to be associated with both promotion and regulation of transcription [4]. For example, mark H3K36me3 (trimethylation of lysine 36 on histone H3) has been correlated with active transcription independent of nucleotide composition [5]. While the mechanism of increasing expression has not been fully investigated, it is believed that the feature may slow RNAPII, which ensures exon inclusion and increases the amount of gene product [5]. Mark H3K4me3 (trimethylation of lysine 4 on histone H3) has been associated with regulation of transcriptionally active genes [6].

Histone modifications have been shown to form a “histone code” with various regulatory functions [7], [8]. These combinations of histone marks can be summarized as “chromatin states.” Specifically, chromatin states are determined through a holistic evaluation of the combination of chromatin marks present in a region. Multiple different methods are available for predicting chromatin states, such as ChromHMM [9], Segway [10], and HMMSeg [11].

Previous work has shown the potential of using several histone modifications to predict levels of gene expression in nearby genes [12]. Integrating information across multiple histone modifications also amplifies the accuracy of predictions, as combinations of marks have been shown to be extra informative [13]. Additionally, next-generation genome sequencing technologies developed in the last twenty years have made it possible to conduct studies encompassing the entire genome rather than just a minute part [14]. Due to the incredible volume of data needed to generate an accurate prediction, machine learning models have become important to revealing biologically significant results.

Here, we combine the power of histone marks and predictive models. We built machine learning models that take predictors (histone modification levels) as input and return measures of gene expression (mRNA levels) as output. Given previous success in similar predictions, we based our method on a two-step model and applied it to a new set of data [15]. Two categories of computer models were utilized in this project: random forest models and linear models. Both have previously been used in the field to associate relationships between predictors and measured expression [16], [4], but to our knowledge have not been used to predict in primary tissues samples, which we included in our dataset. Furthermore, we investigated the relationship between chromatin states and histone mark signals by calculating the fold-enrichment of each state in localized regions.

## Methods

### Overview

The first step to building our predictive models was to localize the histone mark to a relevant region for each sample. For this, we used protein-coding genes longer than 4100 base pairs divided it into 81 “bins,” as shown in Fig. 1. For each epigenome and each histone modification, we localized the signal to the “best bin” where histone mark presence most closely correlated with gene expression, as shown in Fig. 2. Next, we trained our two-step model using half our data as the training set: the model first classified genes as either expressed or unexpressed and then predicted the magnitude of expression for expressed genes using linear regression. Once we had created a predictive model for every sample, we calculated predictive accuracy based on the test set (the remaining half of our data). We averaged our results across sample type and anatomy to observe trends in accuracy and histone mark importance. To further our understanding of chromatin states, we calculated the chromatin state enrichment in each histone mark’s best bin across all samples.

**Figure 1:**
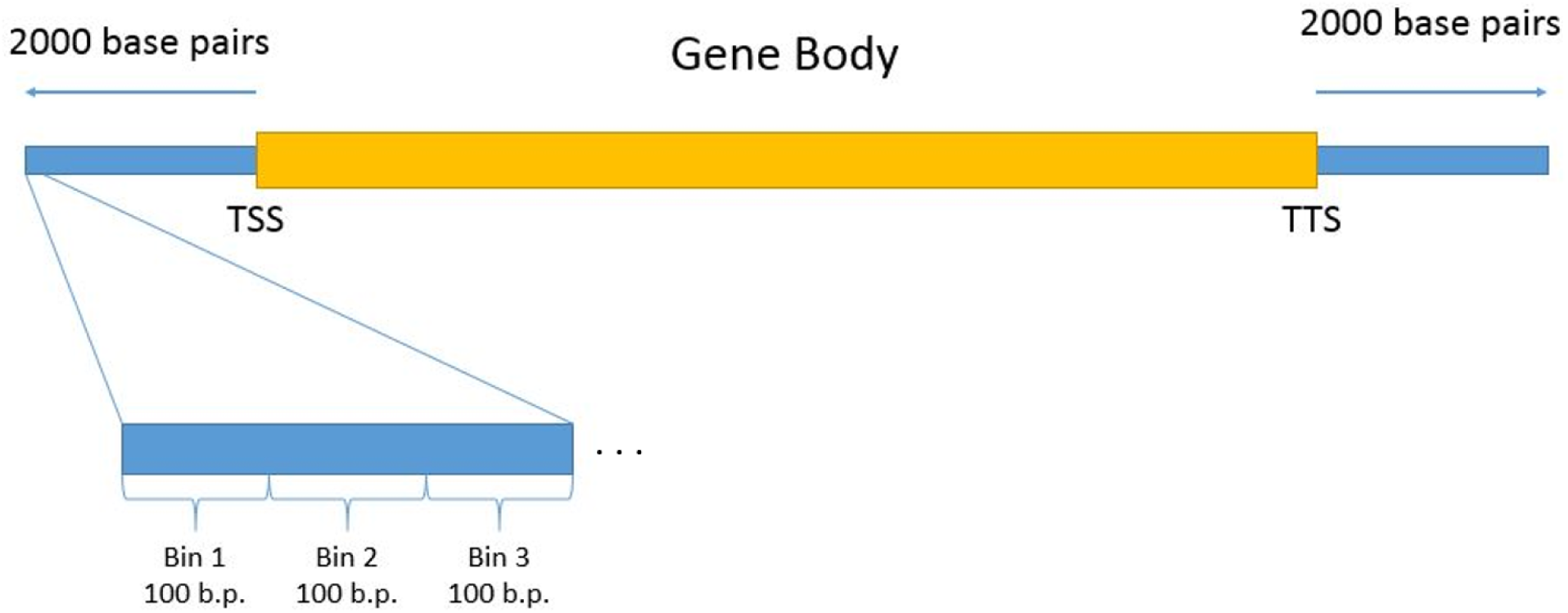
Here, the gene body as well as 2000 base pairs upstream and downstream is divided into 80 bins of 100 base pairs each and one bin to capture the rest of the gene body.

**Figure 2:**
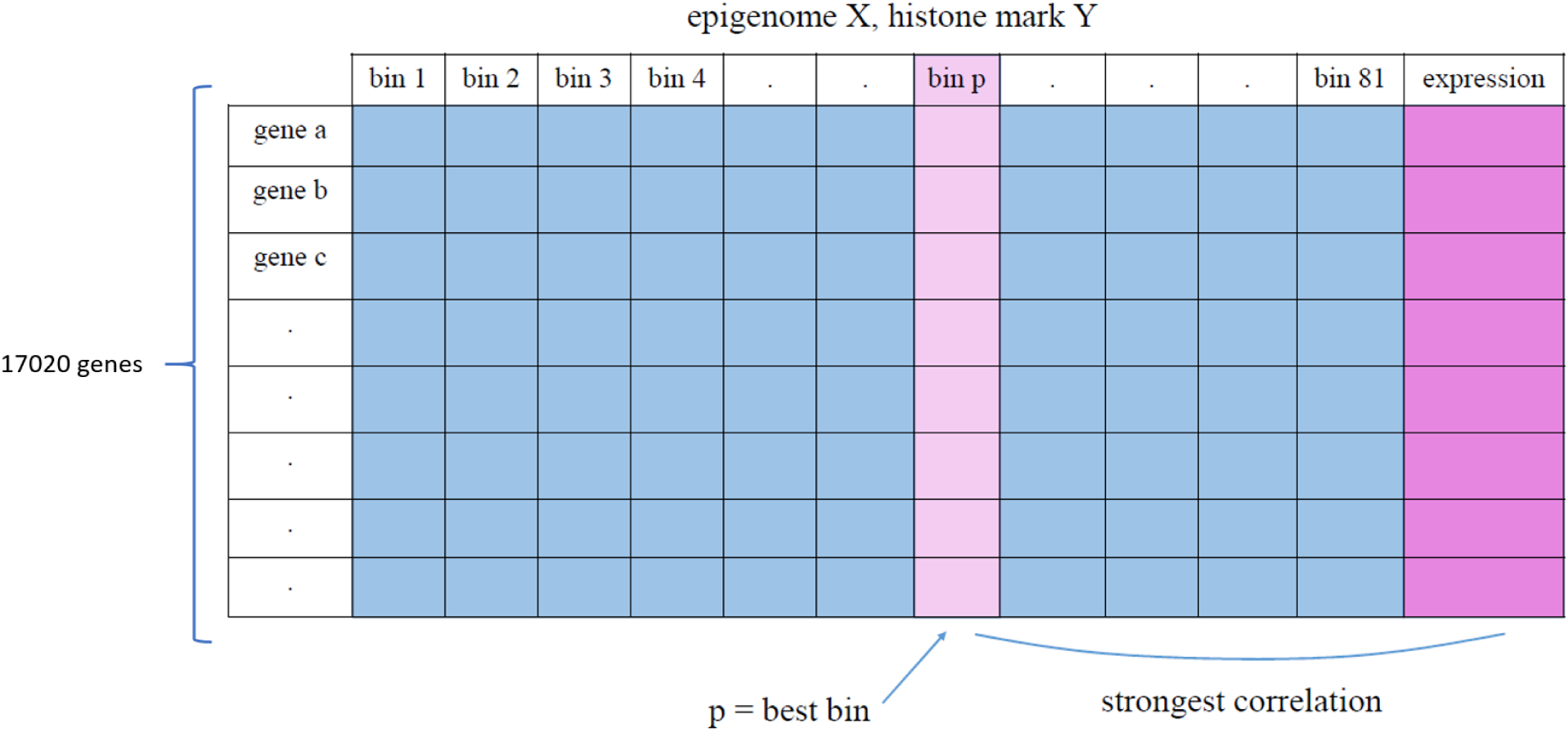
Here, bin p is chosen as the “best bin” for epigenome X and histone mark Y.

### Pre-processing gene information

We began by filtering the GENCODE Ensembl file for unique protein-coding genes longer than 4100 base pairs [17]. The filtered file contained chromosome identification, transcription start and termination site information, an ID, and strand information. Previous work suggested that 4100 base pairs was an appropriate cutoff to reduce the amount of noise that would confuse the models [4], [12].

### Binning genes

As described in [18] and shown in Fig. 1, each gene body (and its flanking 2000 base pairs) was then divided into 81 “bins”: 80 bins of 100 base pairs each and one bin to capture remaining gene body. The file detailing the binned genes and their chromosome, TSS, TTS, ID, and strand information was written in a bed format.

### Measuring histone levels

We used data that recorded the relative presence of histone marks H3K27me3, H3K36me3, H3K4me1, H3K4me3, and H3K9me3 as a continuous, consolidated signal across the genome. With this data, we calculated the relative presence of each histone mark in each of the 81 bins for each epigenome with documented gene expression data using bigWigAverageOverBed [19].

### Best-bin approach

Since histone marks are known to perform different functions in different contexts [13], we chose to localize each histone mark signal to the most relevant relative region of the gene. A “best bin” was selected for each of the five histone modifications for each epigenome, based on the bins described in the section “Binning genes.” As shown in [15] and Fig. 2, the bin in which (across all genes) histone information correlated best with expression data for that epigenome was selected. For each epigenome, the data for the best bin for each histone mark was compiled. The final result (for each epigenome) was a matrix representing the presence of each histone mark and expression data for every gene along the genome.

### Prediction

Due to the massive volume of data required to predict gene expression from histone information, we utilized machine learning methods to build predictive models. Since testing the accuracy of the models on the same data we used to train them would produce artificially high accuracy values, we randomly selected half of the genes to use as the training set and saved the remaining half of the genes for testing. During the process of fine-tuning our models we also implemented cross-validation, but ultimately decided against using it in our procedure given that cross-validation does not produce a final model with linear coefficients to represent relative histone importance.

As described in [15], we used a two-step model to account for the fact that different modifications may control whether a gene is expressed at all and if so, how much it is expressed. For the first step, the random forest classification model (built using the randomForest package in R [20]) classified the genes as “on” (expressed) or “off” (unexpressed) based on whether or not any expression was occurring at said gene [16]. The second step used a linear regression model (built using the lm function in base R [21]) to predict the magnitude of expression for all “on” genes [4]. The regression models produced linear coefficients for each of the five histone marks, which we later used to determine predictor importance. We took the absolute value of each linear coefficient so the value would reflect only the magnitude of importance, regardless of directionality. The model also produced p-values, which we later used to determine whether or not each predictor is significant.

### Summarization

To better observe general trends in our results, we scaled the sum of the linear coefficients to the *r*^2^ value so they would represent both relative importance and accuracy of the model. Specifically, let *c_x_,_a_* be the raw linear coefficient calculated for histone mark x by the model generated for epigenome a and 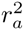 be the *r*^2^ value for the model predicting for epigenome a. Then, to calculate *s_x a_*, the resulting scaled linear coefficient for histone mark x and epigenome a:

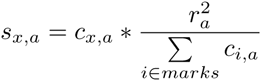

We averaged the scaled linear coefficients recorded for each of the five histone marks in epigenome groups organized by type (as shown in Fig. 3) and organized by anatomy (as shown in Fig. 4). As used in [22], we calculated the standard error of the mean across each of the groups.

**Figure 3:**
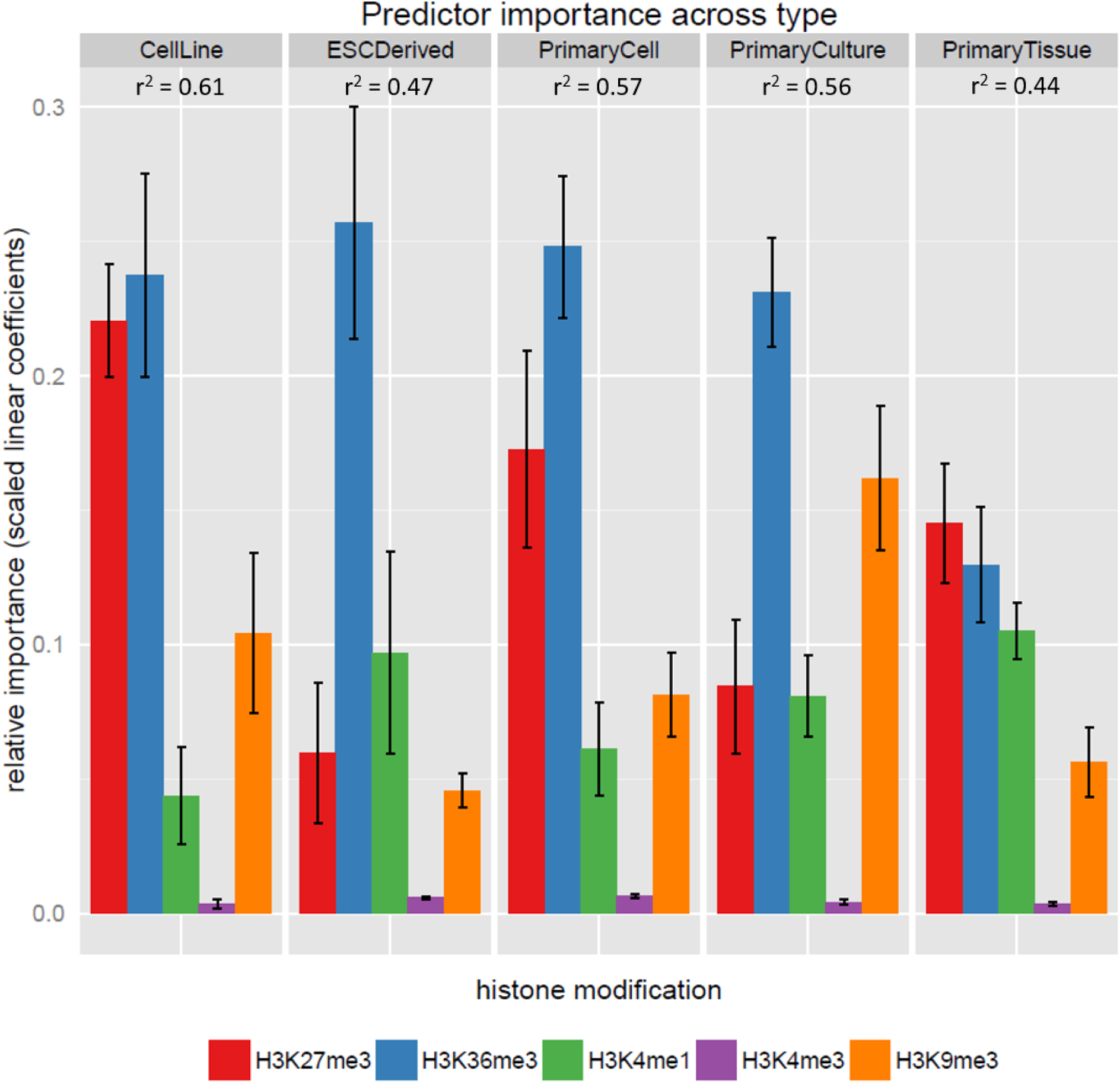
H3K36me3 is the most predictive histone mark in cultured genomes. The height of the bars represents how much weight the model gave the corresponding predictor and are scaled to the *r*^2^ value of that model. Error bars represent standard error of the mean across the category.

Since the *r*^2^ values represented the cumulative accuracy of both steps of the two-step model, we also evaluated the accuracy of the classification step alone by calculating the area under the Receiver Operating Characteristic (ROC) curve, or AUC value, which represents the specificity and sensitivity of the classification.

### Chromatin Annotations

To further our understanding of the relationship between chromatin states and predictive histone mark regions, we also identified which chromatin states were significantly enriched in the best bins we used for prediction. After classifying each bin by whichever chromatin state the majority of its base pairs represented, we identified the background distribution by calculating the average percent occurrence of each chromatin state across all epigenomes. We then looked only at the best bin chosen for each of the five histone marks and calculated the average percent occurrence of each chromatin state within the chosen best bins across all epigenomes. To find the fold-enrichment we took the ratio of the occurrence in each epigenome and the background distribution. We then averaged the fold-enrichment across all epigenomes. Formally, let *N* be the number of genes in each epigenome, *p_x,r,a_* be the number of occurrences of state x in the best bin of histone mark r across epigenome a, and *q_x_* be the average number of occurrences of state x per epigenome (across all bins and all samples). Then, to calculate *f_x,r,a_*, the fold-enrichment of state x in the best bin of histone mark r across epigenome a:

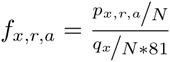

Lastly, we calculated a 95% confidence interval of our data using the standard error of the mean (assuming a normal distribution). Specifically, let *SE* be the standard error of the mean found for enrichment of the specified state and histone mark. Then, to calculate *c*, the 95% confidence interval:

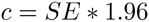

## Results

### Predictive accuracy varies across different sample types

To quantify the accuracy of our predictive model, we calculated the correlation between gene expression predicted by our models and measured gene expression across epigenomes. *r*^2^ values were distributed between 0.35 and 0.65 (see Supplemental Fig. 1), with a mean *r*^2^ value of 0.51 across the 54 samples and a standard deviation of .079.

Furthermore, we looked at predictive ability averaged across epigenome groups based on sample type. Generally, models were most accurate in prediction for cell lines (see Fig. 3), with an average *r*^2^ value of 0.61, and worst for primary tissues, with an average *r*^2^ value of 0.44. This was perhaps due to the heterogeneous nature of the samples classified as primary tissues (which may consist of cells that perform a variety of functions). Although cell lines do not perfectly model complex cells found in the human body, they are often used in the field [15]; our findings suggest a strong relationship between epigenetic marks and gene expression in cell lines, although the extent to which this reflects the state of complex organisms in-vivo is undetermined. Primary cell samples and primary culture samples were predicted with more accuracy than the primary tissues, though not as accurately as the cell lines, with an average *r*^2^ value of 0.56 for both types. ESC derived samples were predicted less accurately, with an average *r*^2^ value of 0.46. This suggests that predictive ability suffers as differentiation of cell types achieves cell specification.

Accuracy of the predictive models varied across sample anatomy as well (see Fig. 4), with average *r*^2^ values ranging from 0.37 to 0.65. Notably, blood and skin samples were predicted with above average accuracy, with average *r*^2^ values of 0.59 and 0.57 respectively. Predictive accuracy of intestine and vascular samples suffered, with average *r*^2^ values of 0.44 and 0.45 respectively, suggesting the need for more complex models to accurately capture the transcriptional state of these samples.

**Figure 4:**
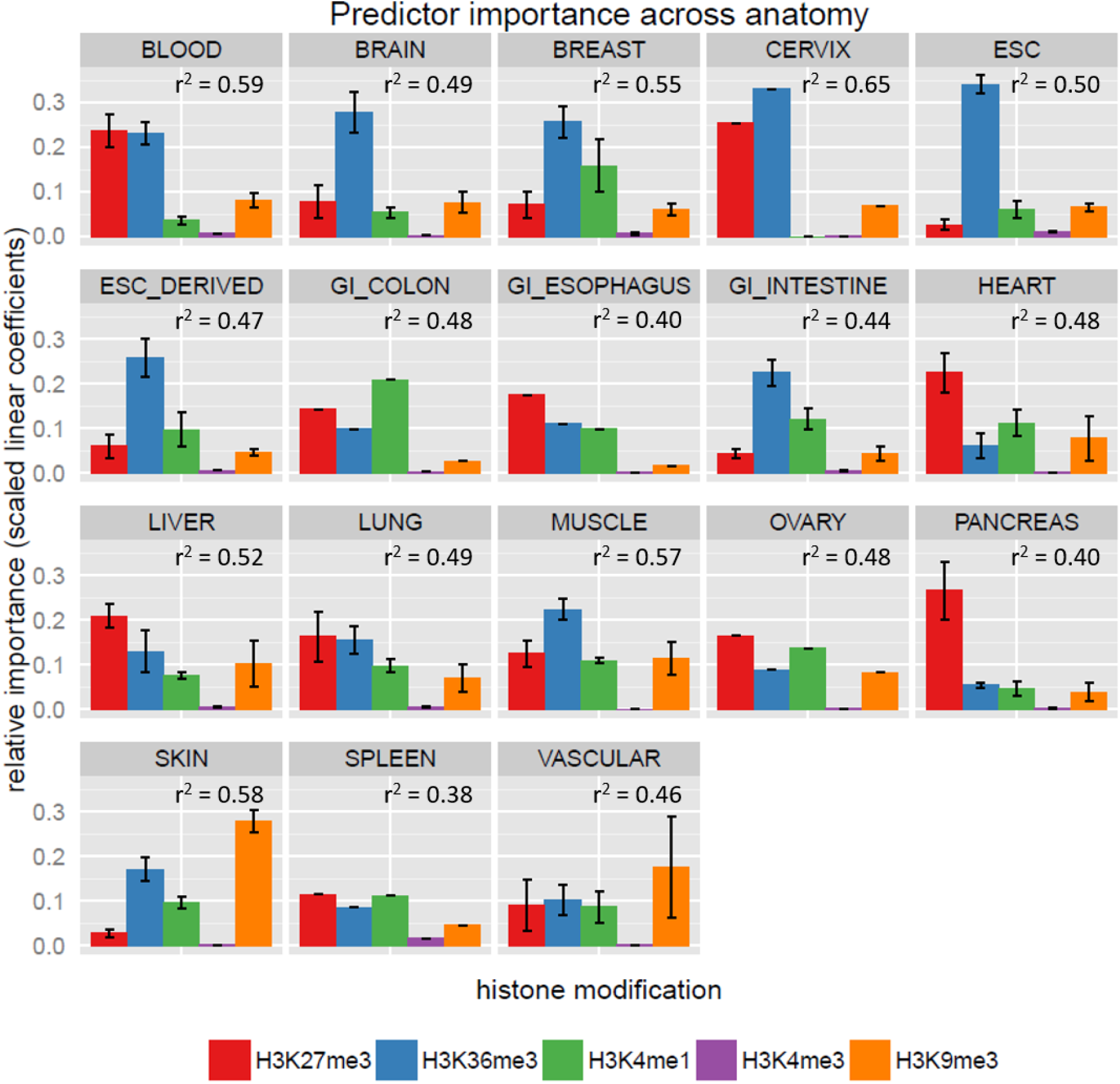
H3K36me3 is the most predictive histone mark across most classes of anatomy. The height of the bars represents how much weight the model gave the corresponding predictor and are scaled to the *r*^2^ value of that model. Error bars represent standard error of the mean across the category.

Our two-step model separately performed classification of genes into expressed and unexpressed categories and secondly regression to determine the magnitude of expression. While the *r*^2^ value quantifies accuracy of the model as a whole, we can also study the accuracy of just the classification step through the Receiver Operating Characteristic (ROC) curve. By balancing sensitivity with specificity, the area under the ROC curve (AUC) value quantifies classification accuracy. For example, we can achieve a sensitivity of 0.87 with a specificity of 0.89 for epigenome E054. We found that our models generally achieved high AUC values that were consistent across all epigenomes. Specifically, the mean AUC was 0.91 with a standard deviation of 0.018. AUC values reached a maximum of 0.94 (see Supplemental Fig. 5).

### Relative weight of histone marks varies across different anatomical groups

Our models were built on data collected by monitoring levels of histone marks H3K36me3, H3K27me3, H3K4me1, H3K9me3, and H3K4me3. Over all epigenomes, we calculated the relative importance of each histone mark as the sum of the scaled linear coefficients (see Methods for scaling procedure) assigned to them by all linear models (see Supplemental Fig. 2). We found that mark H3K36me3 was the most important mark by far (relative weight = 0.394), affirming previous results [5], [15]. H3K4me3 was given very little weight by the models (relative weight = 0.00875), while the remaining three marks were all weighted approximately equally (relative weight of H3K27me3 = 0.235, relative weight of H34Kme1 = 0.171, relative weight of H3K9me3 = 0.191).

Relative importance was not completely consistent across sample types (see Fig. 3). We found that mark H3K36me3 was only given a significantly higher weight in three of the five cell types we studied (ESC derived, Primary Cell, Primary Culture). In the remaining two sample types, H3K36me3 was not significantly more important than mark H3K27me3. Relative weight varied across sample anatomy as well (see Fig. 4). Across anatomical groups with more than one sample, H3K36me3 was significantly the most important mark in six of the 13 groups (brain, breast, ESC, ESC derived, intestine, muscle), H3K27me3 was significantly the most important in three (heart, liver, pancreas), and H3K9me3 was significantly the most important in one (skin).

While linear coefficients allowed us to approximate importance of each mark, we also wanted to identify the statistical significance of each mark in our model. To do this we calculated the p-values associated with the linear regression; specifically, the statistical likelihood that the true coefficient for each mark was not 0. We found that the relative impact (effect size) of each mark was not always correlated with the statistical significance of that mark having an effect (see Supplemental Fig. 2). For example, we found that the H3K4me3 (very unimportant), H3K27me3 (relatively important), and H3K4me1 (relatively important) were significant for more than 98% of the epigenomes, while H3K36me3 (very important) was significant for 93% of the epigenomes and H3K9me3 (relatively important) was significant for 73% of the epigenomes. These results confirm previous work which suggests that histone marks have a statistically significant relationship with gene expression, although future work will have to be done to discern whether the relationship is causative or merely correlative. Further, our findings suggest that when prioritizing histone marks for prediction of gene expression, scientists will have to balance the effect size with the statistical significance of the histone mark in choosing the best combinations of marks.

### Chromatin state enrichment varies across localized regions

We identified chromatin state enrichments and depletions among the best bins chosen for the five histone marks we studied in each sample (see Fig. 5). We used a 95% confidence interval to determine whether enrichments were signficant (see Methods for confidence interval calculation). Overall, TssA and TssBiv were the most frequently enriched states, significantly enriched in best bins for three of the five histone marks (H3K27me3, H3K4me3, H3K9me3); this makes sense as active promoters are known to play a strong regulatory role in gene expression [23]. TxWk was the one of the second most frequently enriched state, significantly enriched in best bins for two of the five histone marks (H3K36me3, HeK4me1). Strikingly, TssA/TssBiv and TxWk were enriched in the best bins of mutually exclusive marks, which perhaps reflects a tendency of H3K27me3, H3K4me3, and H3K9me3 to be localized near the transcription start site (TSS), while H3K36me3 and H3K4me1 were often localized near the gene body.

**Figure 5:**
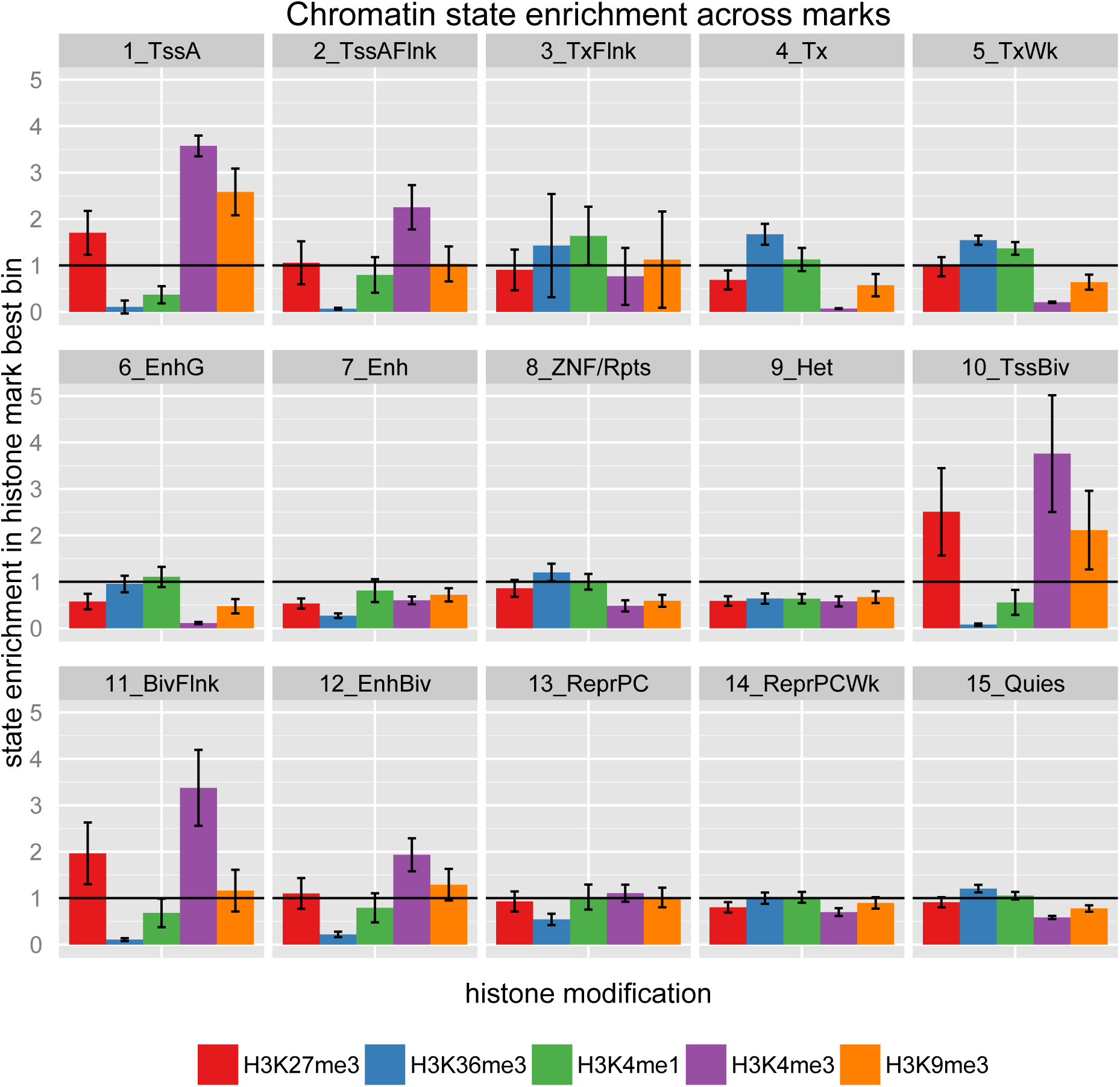
TssA, TssBiv, and TxWk are the most frequently significantly enriched states. The height of the bars represents the fold-enrichment of the represented state in the best bin of the represented histone mark. Error bars represent 95% confidence intervals.

At least one state was significantly enriched in for every histone mark, though some had a higher number of significant chromatin state enrichments than others (see Fig. 6). For example, among the best bins chosen for mark H3K4me3, five states were significantly enriched (TssA, TssAFlnk, TssBiv, BivFlnk, EnhBiv), and among the best bins chosen for mark H3K36me3, four states were significantly enriched (Tx, TxWk, ZNF/Rpts, Quies). Mathematically, the presence of enrichments necessitates depletion of other states. Interestingly, among the best bins chosen for mark H3K36me3, the significantly depleted states include the same five states that were enriched for mark H3K4me3 (TssA, TssAFlnk, TssBiv, BivFlnk, EnhBiv) as well as Enh and Het. Among the best bins chosen for mark H3K4me3, the significantly depleted states include the four states enriched for mark H3K36me3 (Tx, TxWk, ZNF/Rpts, Quies), as well as Enh and Het, which were also significantly depleted for mark H3K36me3. Notably, we find significant enrichments for states TxFlnk and TxWk among the best bins chosen for mark H3K4me1, which likely reflects the strong presence of H3K4me1 in the profile for those states. In this analysis, we do not disentangle the role of the histone marks in identifying chromatin state when calculating these significant correlations. However, our preliminary results suggest that further study of the relationship between localized histone mark regions and chromatin states could be fruitful.

**Figure 6:**
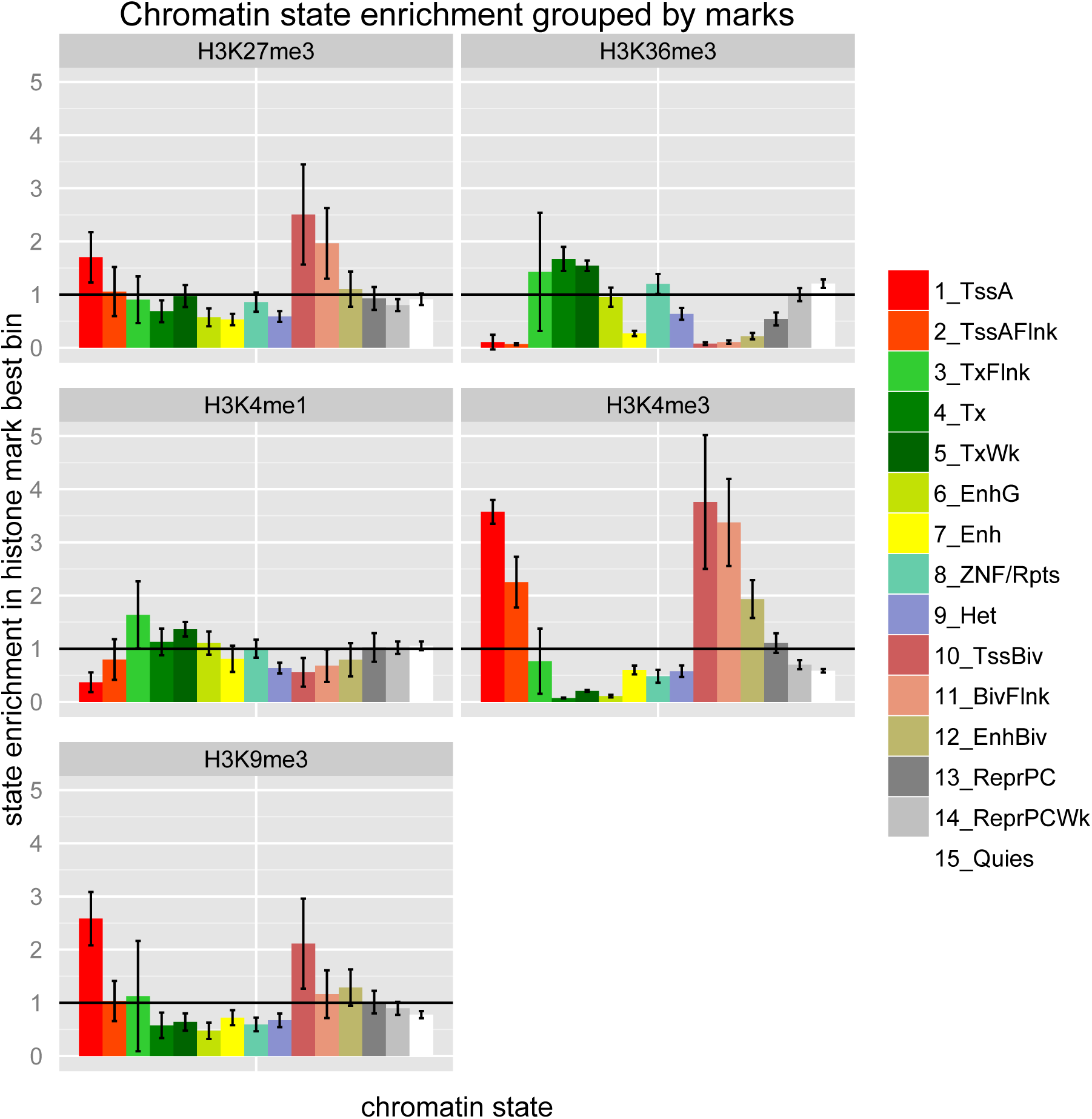
Marks H3K36me3 and H3K4me3 contain the most significantly enriched states. The height of the bars represents the fold-enrichment of the represented state in the best bin of the represented histone mark. Error bars represent 95% confidence intervals.

### Discussion

These findings have multiple implications for the field as a whole. The range of predictive accuracy we achieved across sample types suggests that tissues are more difficult to predict using machine learning algorithms, and therefore more complex regulatory models are needed to attain the same level of accuracy as in cultured genomes. This makes sense as tissue samples are often more heterogeneous than cell lines due to cell differentiation and specification.

Biologically, our work supports the finding that mark H3K36me3 is heavily involved in positive regulation (promotion) of gene expression. The differences we found across sample type and anatomy suggest that histone mark regulation is sample-dependent, which can inform future research to study differences in tissue-specific regulation. From a more practical standpoint, this can inform choices researchers are financially forced to make about what chromatin features and samples to study that will yield productive results, given limited budgets.

Our preliminary results regarding chromatin states reveal an underlying relationship with histone modifications and gene expression. This suggests a potential for predictive models for expression based on chromatin states, rather than individual histone marks. It also provides a new view for chromatin state histone mark profiles narrowed to the localized regions which are most relevant for each histone modification.

Our results bring up several interesting questions. While we used the default parameters of the random forest classification model in R, future work could be done to optimize the sensitivity and specificity of the classification step, depending on the priorities of the predictive model. Investigating the ROC (Receiver Operating Characteristic) curve could determine the best cutoffs to improve predictive power. Furthermore, future predictive models could more precisely model promoter regions, as our best bin procedure only looks 2000 base pairs upstream and downstream of the gene body. Lastly, the importance of distant regulatory regions should not be overlooked. Future models that incorporate analysis of relevant distant genomic regions would likely enhance the accuracy of our models and provide more information about the relationship between histone marks and gene expression.

These results provide novel insights into epgienetic regulation of transcription in contexts which have not previously been extensively studied. The breadth of our data set, including five different sample types, and unusual localization procedure both validate previous work which has been done in the field as well as suggesting new avenues for productive research.

## Supplemental figures

**Supplemental Figure 1:**
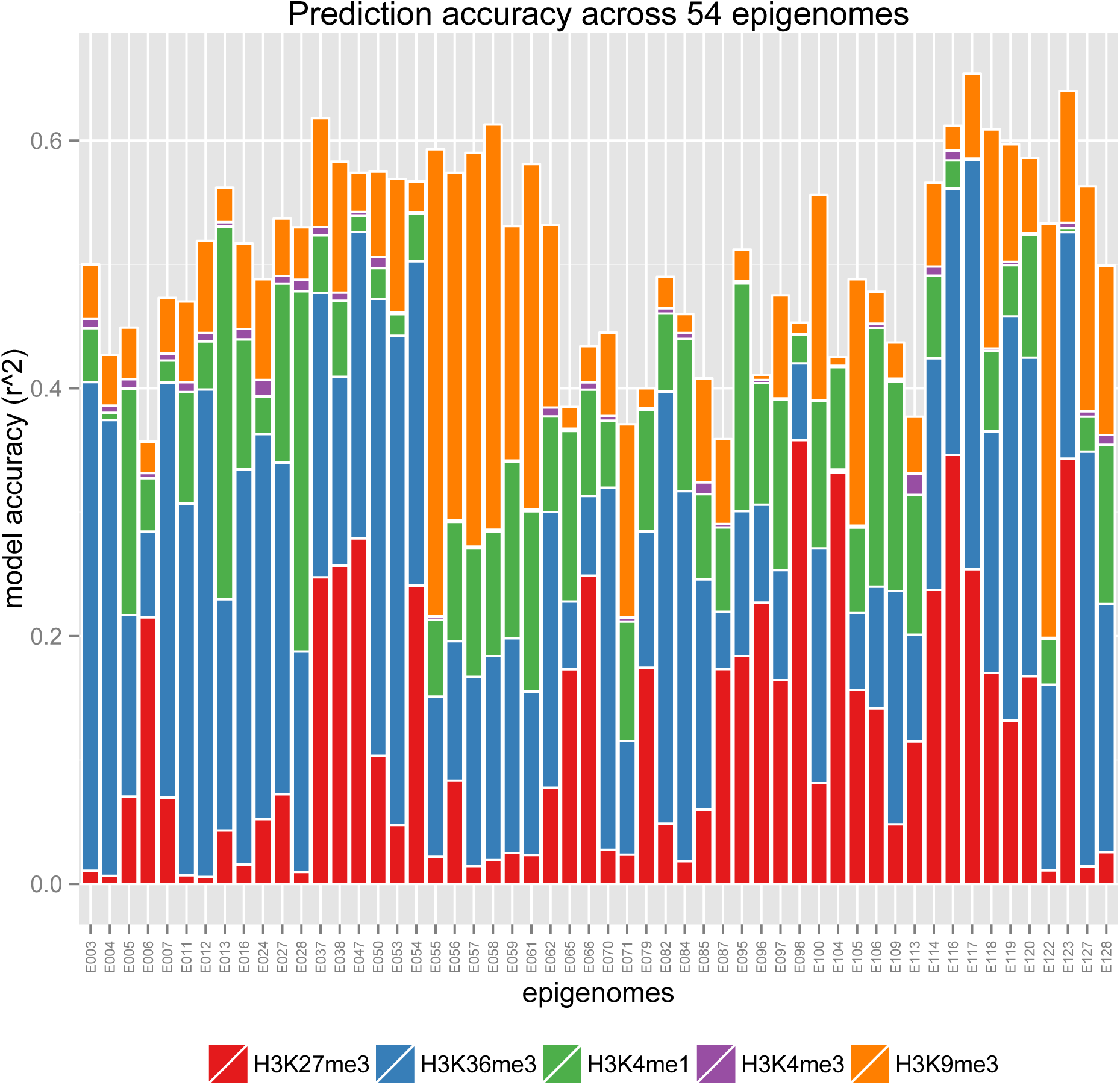
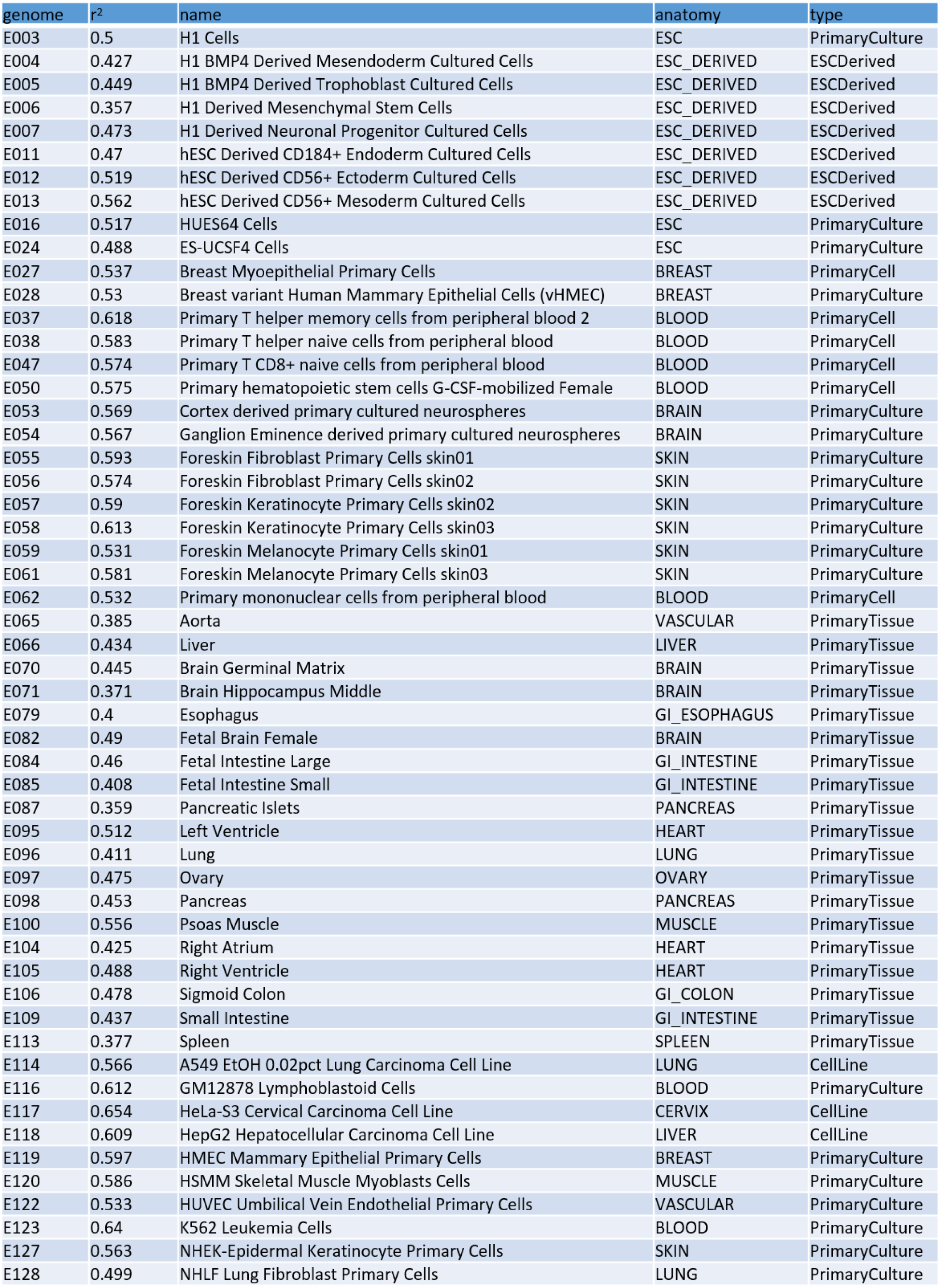
*r*^2^ values are evenly distributed between 0.35 and 0.65. The height of the bars represents the *r*^2^ value of that model and the height of each colored section represents how much weight the model gave the corresponding predictor.

**Supplemental Figure 2:**
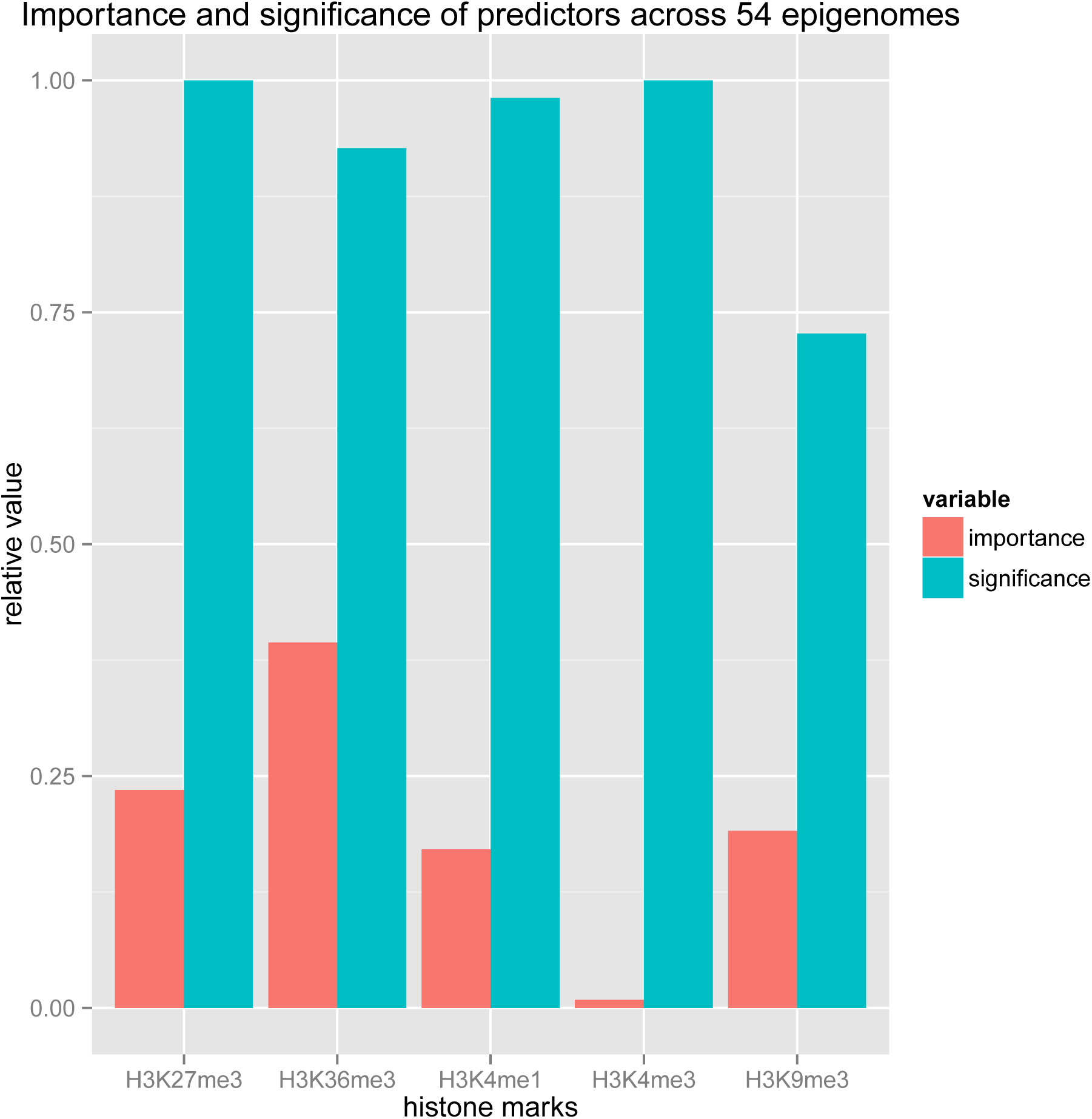
H3K36me3 is overall the most relatively important histone mark, but all 5 marks have a high level of significance. The height of the “importance” bars represents the relative weight of the histone mark (see Results for calculation of the relative weight) and the height of the “significance” bars represents the fraction of samples in which the histone mark was significant in prediction.

**Supplemental Figure 3:**
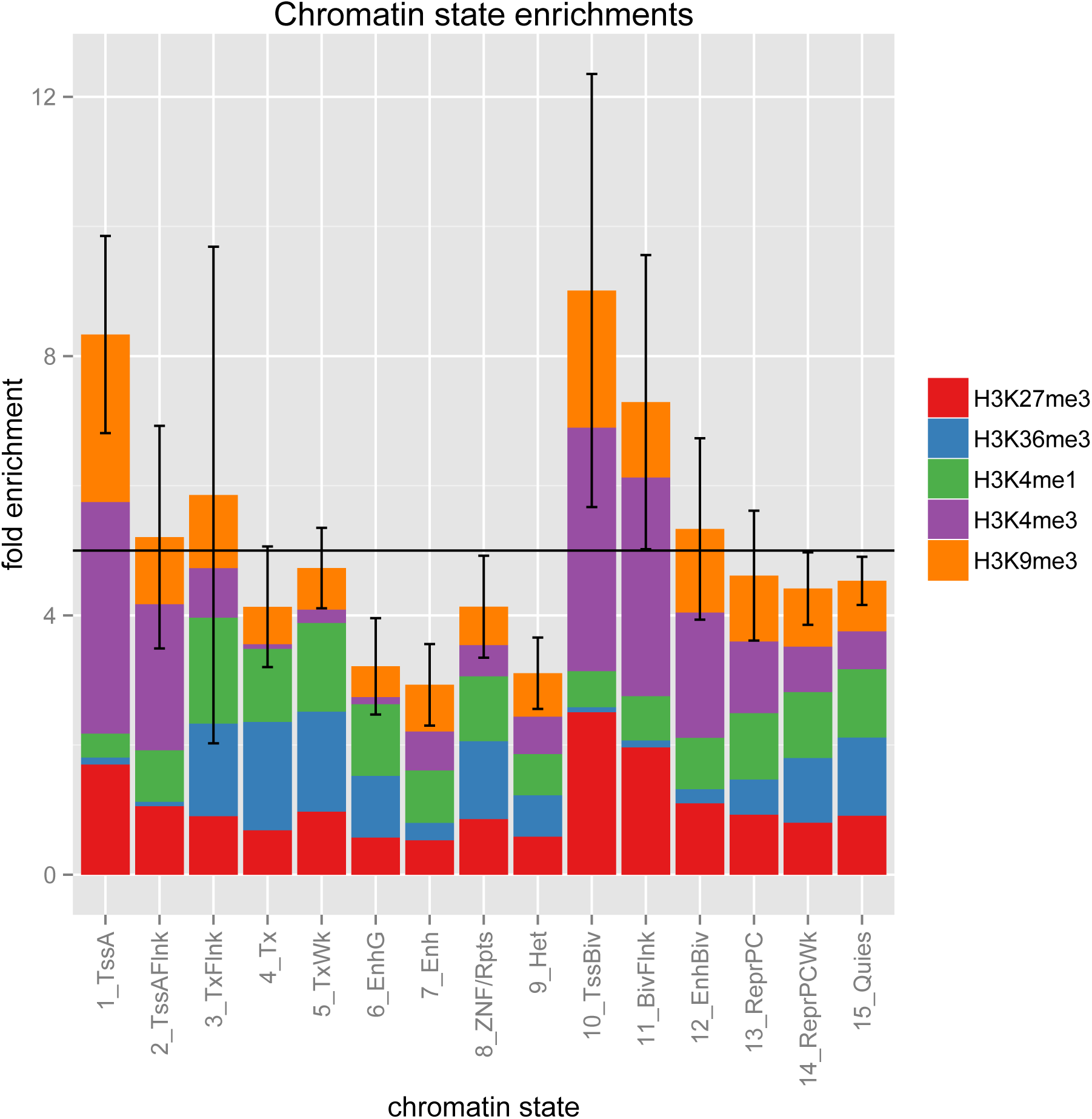
TssA, TssBiv, and BivFlnk are overall the most enriched states. The height of the bars represents the fold-enrichment of the represented state and the height of each colored section represents the fold-enrichment in the best bin of the corresponding histone mark.

**Supplemental Figure 4:**
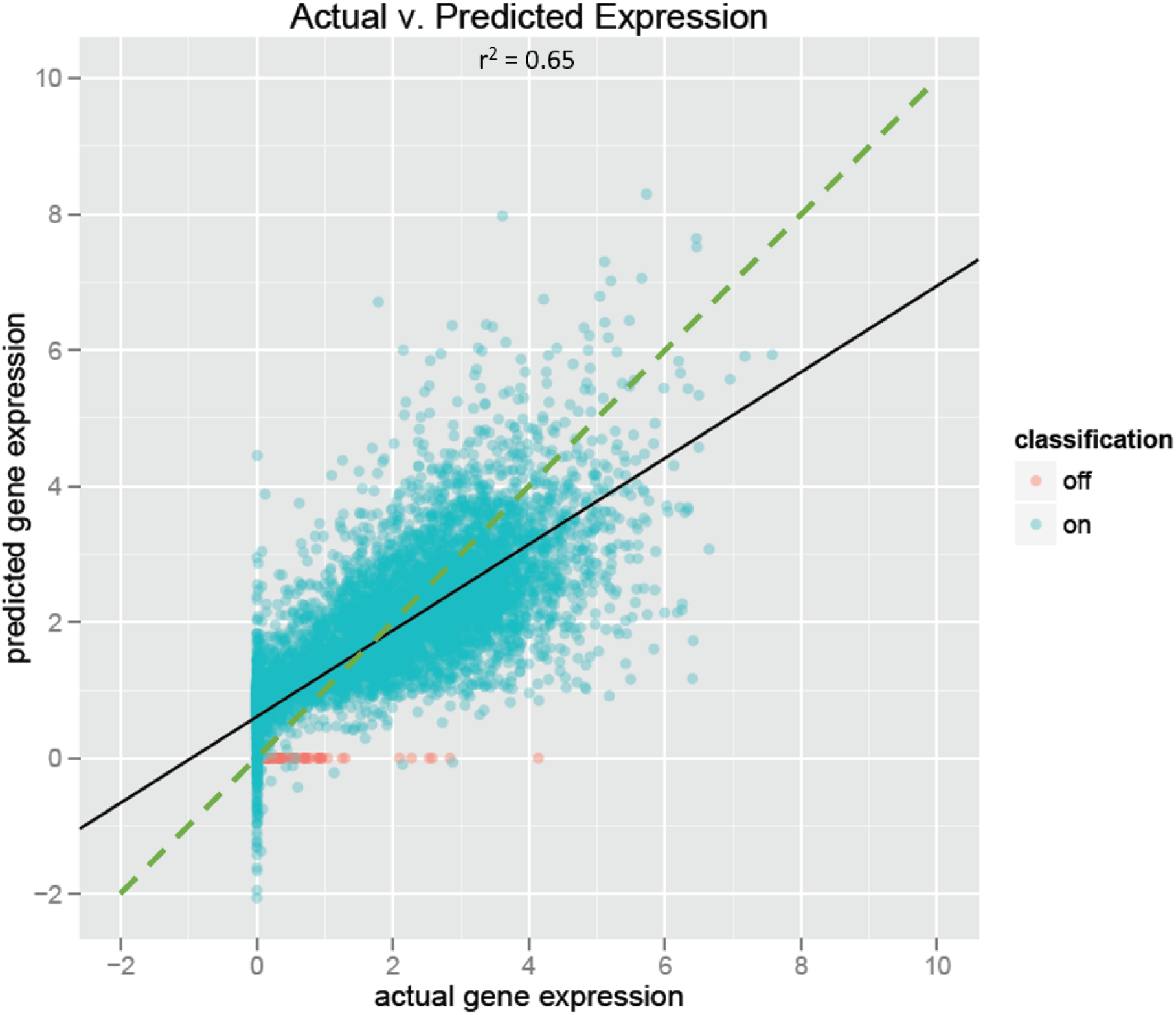
Actual vs. predicted gene expression for HeLa-S3 Cervical Carcinoma Cell Line (highest *r*^2^ value). The solid black line represents the best-fit line for this model, while the dotted green line (y = x) represents perfect prediction.

**Supplemental Figure 5:**
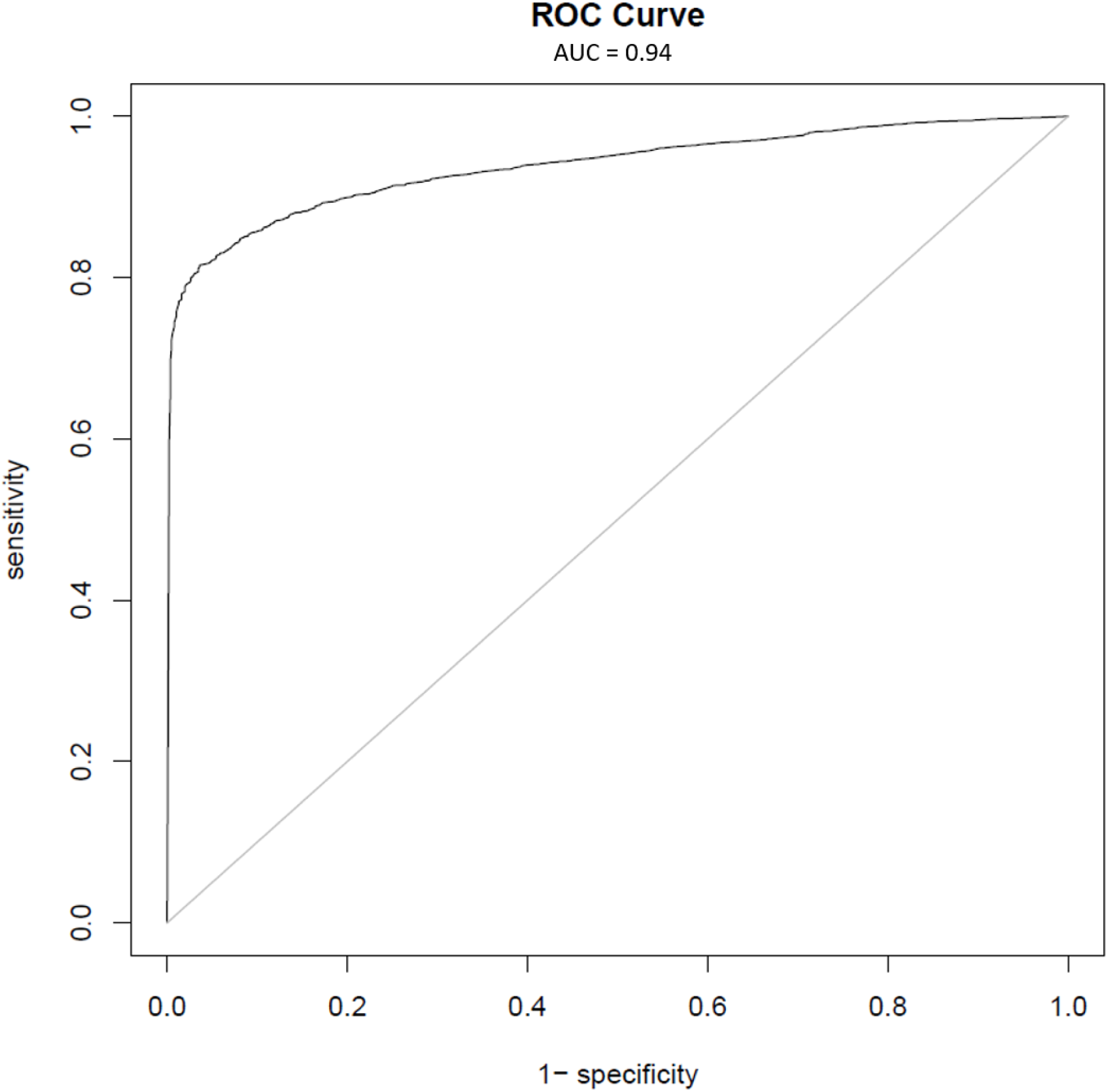
ROC curve for Ganglion Eminence derived primary cultured neurospheres (highest AUC value).

